# Mesenchymal Stem Cell Chondroinduction on Cellulose-Silk Composites is Driven by Substrate Elasticity

**DOI:** 10.1101/383307

**Authors:** Runa Begum, Wael Kafienah

## Abstract

Biomaterials that can physically control the fate of stem cells are critical for the application of *in situ* tissue engineering. However, the mechanisms underlying this phenomenon remain unclear. In this study, cellulose and silk composite substrates, known to induce physically the chondrogenic differentiation of human mesenchymal stem cells (MSCs), were regenerated to fabricate two-dimensional film surfaces. MSCs were grown on these surfaces in the presence of chemicals that interfere with the biochemical and mechanical signalling pathwys of MSCs. The data shows that preventing substrate surface elasticity transmission results in significant downregulation of chondrogenic gene expression. Interference with the classical, chondrogenic Smad2/3 Phosphorylation pathway does not impact the chondrogenic capacity of the substrates. The results highlight the importance of the substrate mechanical elasticity on chondrogenic MSCs and its independence of known chondrogenic biochemical pathways.

## Introduction

Smart biomaterials with inherent capacity to control stem cell can have significant impact on the field of cartilage tissue regeneration. Currently, the majority of studies demonstrating the successful use of biomaterials for cartilage tissue engineering require the addition of specific growth factors to induce chondrogenic differentiation and control stem cell fate [1]–[4], which are usually animal derived and costly. Eliminating the need for exogenous biological stimulation would not only reduce the costs of production, but also circumvents the biological adverse effects associated with such biologics; for example, intra-articular injections of transforming growth factor-β (TGF-β) results in osteophyte formation in the murine joint [5].

The ideal biomaterial for cartilage tissue engineering would provide a number of attributes that closely align to that of the stem cell niche, which plays a critical role in maintaining stem cell function and fate through orchestrating a plethora of physical and chemical cues to regulate cell proliferation, differentiation, migration and apoptosis [6] [7]. For example, the regulation of localised concentrations of TGF-β is essential, as TGF-β is a potent agonist for chondrogenic differentiation [8]. Binding of this growth factor to the type II receptor (TBR-II) initiates the signalling pathway leading to the recruitment of a type I receptor (TBR-I) followed by the downstream phosphorylation of Smad2 and Smad3 [9]. These Smad proteins then associate with the co-Smad, Smad4, forming a protein complex, resulting in its translocation to the nucleus where it can regulate transcriptional activation. The principal transcription factor regulating chondrogenesis is the SRY-type HMG box 9 (Sox9), activation of which, in combination with TGF-β signalling, is key for chondrogenic differentiation [10] [11]. Sox9 activates its target gene Col2a1 which encodes type II collagen, the main cartilage extracellular matrix (ECM) protein. This transcription factor is also involved in the upregulation of aggrecan, another essential cartilage ECM protein [12].

Mechanical aspects of a cell external environment may also influence its fate. For example, biomaterial elasticity has been identified as a driving factor in determining MSC lineage specification. Engler *et al*. demonstrated that MSCs differentiate toward the lineage respective of the substrate elasticity upon which they are cultured [13]. Soft matrices mimicking that of brain led to cells differentiating into a neuronal phenotype; cells seeded on stiffer matrices, as that seen in muscle displayed a myogenic phenotype; and that on rigid matrices, as in collagenous bone, became osteogenic. In an effort to understand this complex cell-biomaterial dependency, Engler *et al*. showed that addition of Blebbistatin during cell culture prevented force transmission *via* cell surface focal adhesions, and therefore prevented elasticity-driven differentiation of the MSCs [13]. Blebbistatin is a potent inhibitor of nonmuscle myosin II (NMM II) [14], a component of the cell actin cytoskeleton important in force transmission between intra- and extra-cellular environments.

Beyond surface elasticity, a number of other scaffold-dependent morphological and chemical properties can influence stem cell differentiation. Addressable chemical functional groups on surfaces interacting with MSCs can direct their differentiation down specific lineages [15], perhaps through the activation of specific differentiation signalling pathways. Curran *et al*. showed that when MSCs were seeded on silane-treated glass surfaces functionalised with hydroxyl or amide groups, the cells expressed chondrogenic and osteogenic mRNA respectively, in the absence of stimulating factors [15]. The nanoscale topography of a biomaterial may also influence stem cell fate. Dalby *et al*. demonstrated how the structural organisation of an *in vitro* matrix can influence MSC differentiation [16]. Cells grown on semi-disordered substrates expressed calcifying bone proteins, whereas those seeded on flat substrates showed no such induction.

We have shown that composites of the natural polymers cellulose and silk can induce chondrogenic differentiation of human MSCs [17]. The neat and composite materials differed in their material stiffness (elasticity), tensile strength, nanotopography and surface functional groups [17]. Not only could these natural polymer surfaces support the growth of MSCs, they could drive their chondrogenic differentiation without the addition of soluble stimulating factors. The blend ratio of 75% cellulose and 25% silk upregulated chondrogenic genes and led to the deposition of cartilage extracellular matrix (ECM) proteins, demonstrating the potency of this composite. Surface elasticity and chemical functional groups were identified as the key differential factors between the natural polymer substrates fabricated by Singh *et al*. [17]. However, the molecular mechanisms driving the chondrogenic differentiation seen on these composites remain unknown.

Here we demonstrated that preventing substrate surface elasticity transmission results in significant downregulation of chondrogenic gene expression whilst interference with the chondrogenic Smad2/3 Phosphorylation pathway does not impact the chondrogenic capacity of the substrates. The results highlight the importance of the substrate mechanical elasticity on chondrogenic MSCs and its independence of known chondrogenic biochemical pathways. This information will contribute to better design of chondroinductive biomaterials and allow the development of smart, 3D natural polymer scaffolds pertinent to the clinic.

## Materials and Methods

### Preparation of Cast Films

*Bombyx mori* silk (Aurora Silk, Portland, USA) was degummed to obtain silk fibroin fibres as follows; fibres were cut into 1 cm pieces and boiled in 0.02M Na_2_CO_3_ solution. Fibres were then washed thoroughly with distilled water to ensure removal of sericin proteins and left to dry overnight in a fume hood. Use of silk fibroin shall be referred to as silk from herein. Cellulose from wood pulp, degree of polymerisation (DP) 890, was purchased as sheets (Rayonier Inc., Jacksonville, USA) and ground to a powder. Composite cast films of cellulose:silk in a 75:25 ratio was prepared as outlined by Singh *et al*. [17]. Briefly, cellulose and silk polymers were weighed to prepare a 1.5% polymer conc. (at the appropriate ratio) in 5g of 1-ethyl-3-methylimidazolium acetate (EMIMAc, Sigma Aldrich). The ionic liquid containing the polymers was then heated at 85°C with stirring, for 2hrs. Polymer solutions were then poured into pre-heated glass dishes and left to cool overnight. Following overnight period, ethanol:acetic acid (EtOH:AcOH 90:10 respectively) was added to the composite films to aid coagulation of the polymers. Samples were covered and left overnight. Following the overnight period, any remaining solvent was removed, and films immersed in distilled water. The water was changed regularly (up to twice daily) until all ionic liquid (EMIAc) had been removed (denoted by change of film from light yellow to clear). Once all ionic liquid had been removed, the 2D films were dried overnight on parafilm.

### Characterisation of Cast Films

Scanning electron microscopy (SEM) was used to image the composite substrates using a using a field emission gun scanning electron microscope (Zeiss EVO Series SEM, Carl Zeiss, UK). The materials were placed on a carbon pad and sputter coated with gold in argon, with a plasma current of 18mA for 15 seconds, (SC7620 Mini Sputter Coater, Quorum Technologies, Kent, UK). The resulting coat thickness was 45.3Å. Samples were then imaged at 15kV accelerating voltage and 24 mm working distance.

Fourier transform infrared spectroscopy (Spectrum 100 FTIR spectrometer, PerkinElmer, MA, USA) analysis was performed in transmission mode across a spectral range of 4000 - 600 cm^−1^. Cast composite substrates were analysed as well as the raw polymers to investigate any difference in chemistry following dissolution and regeneration. Spectra were generated four times per sample to ensure chemical homogeneity throughout samples.

### Human Mesenchymal Stem Cell Culture

Mesenchymal stem cells (MSCs) were isolated from bone marrow plugs recovered from patients undergoing complete hip replacement arthroplasty. Sample collection was carried out following local ethical guidelines in Southmead Hospital, North Bristol Trust. MSCs from four patients were used (*n=4*) in the study. Following isolation, MSCs were expanded in stem cell expansion medium consisting of Dulbecco`s modified Eagles medium (DMEM), 10 % (v/v) fetal bovine serum (FBS, Thermo Scientific Hyclone, UK), 1 % (v/v) Glutamax (Sigma, UK), and 10 % (v/v) penicillin (100 units/mL)/streptomycin (100 mg/ml) antibiotic mixture (P/S, Sigma, UK). The stem cell expansion medium was supplemented with 5ng/ml fibroblast growth factor 2 (FGF-2, PeproTech, UK). FGF-2 is known to enhance bone marrow MSC proliferation and differentiation potential [18], [19]. Cells were cultured at a density of 2.0×10^5^ cells per cm^2^ and incubated at 37 °C in a humidified atmosphere of 5 % CO_2_ and 95 % air. Media was changed every 2-3 days and cells were passaged upon reaching 80-90 % confluency. Cells used for all experiments were between passage 2 and 5.

### Preparation of Cast Films for Stem Cell Culture

The composite substrates were sterilised and fibronectin coated following the protocol established by Singh *et al*. [17]. Briefly, cast films were cut into 8 millimetre discs using a biopsy punch (Stiefel, Schuco Intl). Discs were disinfected as follows; aqueous 70 % ethanol (v/v) was added to tissue culture plastic containing discs for 30 minutes and then washed three times with sterile phosphate buffered saline (PBS) to remove any residual alcohol. Following the wash step, PBS was removed and 100 μg/ml human plasma fibronectin in PBS was added (Sigma, UK). Adsorption of human plasma fibronectin on polymer surfaces enhances cell adhesion [20], [21]. Coated scaffolds were then incubated at 37°C in a humidified atmosphere of 5 % CO_2_ and 95 % air overnight. Following overnight incubation, scaffolds were transferred to 24-well ultra-low attachment plates (Corning, Costar, Sigma UK) containing sterile PBS. The PBS was then removed, enabling complete placement of the discs at the bottom of the well. The ultra-low attachment plate wells are coated with an inert hydrogel layer that inhibits cell adhesion (Corning, Costar, Sigma UK). Plates were covered and left at the back of the tissue culture hood to allow the scaffolds to dry (~2 hours). Stem cell expansion media (as described above) was then added to each well and plates transferred to the incubator (37 °C, 5 % CO_2_, 95 % air) until MSCs were ready for loading.

### Cell Loading on Composite Films

Once MSCs had reached 80-90 % confluency, cells were trypsinised and counted. Spent media was removed from the flask and cell surface washed with sterile PBS. Trypsin-EDTA solution (0.5 %, Sigma, UK) was then added and flask incubated (37 °C, 5 % CO_2_, 95 % air) until cells detached from the plastic surface (~5 minutes). An equal volume of stem cell expansion media was then added, and the cell suspension centrifuged at 1500 revolutions per minute for 5 minutes at 20°C. The supernatant was then removed, and the cell pellet resuspended in expansion media. Cells were counted using a haemocytometer and plated at a density of 28×10^3^ cells per cm^2^ on scaffolds and tissue culture plastic controls.

### Gene Expression Studies

Composite cellulose:silk 75:25 substrates were cultured with MSCs in stem cell expansion media supplemented with 10 ng/ml FGF-2, as per the media conditions used previously [17]. Tissue culture plastic controls were also included as follows; MSCs were cultured in standard 24-well tissue culture plates in stem cell expansion media supplemented with 5 ng/ml and 10 ng/ml FGF-2. MSCs were also cultured on plastic in chondrogenic differentiation media as a positive control for chondrogenesis. The chondrogenic differentiation media consisted of high glucose DMEM containing 1 mM sodium pyruvate, 1 % ITS, 1 % P/S, 50 μg/ml ascorbic acid-2-phosphate and 1×10^−7^ M dexamethasone (all Sigma, UK). This was supplemented with 10 ng/ml transforming growth factor-3 (TGF-β3, R&D Systems). A further control was also included – MSCs cultured in chondrogenic differentiation media without TGF-β3. All cell cultures were incubated at 37 °C, 5 % CO_2_, 95 % air and media changed every 2-3 days. In all cases, cells were cultured for 14 days before downstream genetic analysis.

RNA extraction was performed on composite substrate and tissue plastic control cell cultures after 2 weeks using the RNeasy Plus Mini Kit (Qiagen) and PureLink RNA Mini Kit (Thermo Fisher Scientific), respectively, following manufacturer`s instructions. RNA concentration and the purity of elutes was determined spectrometrically at 260 and 280 nm wavelengths (NanoPhotometer P-class Spectrophotometer, GeneFlow, UK). 20 ng/ml of RNA elution was reverse transcribed to produce complementary deoxyribonucleic acid (cDNA) using the High Capacity cDNA Reverse Transcription Kit (Applied Biosystems) according to the manufacturer`s protocol.

Chondrogenic gene expression was quantified using digital polymerase chain reaction (qPCR, StepOne Plus, Life Technologies). qPCR was performed on 96-well plates with each sample in triplicate. Each well contained 1 μl cDNA (of 20 ng/ml concentration), 3.5 μl nuclease-free water, 0.5 μl TaqMan^®^ Gene Assay primer and 5 μl TaqMan^®^ Gene Expression Master Mix. The following TaqMan^®^ Gene Expression Assays were used; Collagen type II alpha 1 (Col2), Aggrecan (Agg), SRY-box 9 (Sox-9), Collagen type I alpha 1 (Col1) and the housekeeping gene actin beta (ACTB), all purchased from Life Technologies Ltd. Data was analysed using the double delta Ct Analysis method. Gene expression levels of Col-2, Agg, Sox-9 and Col-1 were quantified relative to the expression levels of the housekeeping gene ACTB.

### Extracellular Matrix (ECM) Protein Deposition

Following 21 days culture of MSCs on the composite cellulose:silk 75:25 substrate, ECM protein deposition was assessed using immunohistochemistry. Briefly, cells were washed twice with PBS then incubated for 20 minutes in 4 % (w/v) paraformaldehyde at room temperature. Cells were then washed again with PBS (three times) then permeabilised in 0.1 % Triton X-100 (room temp.). Following a further PBS wash step, the samples were incubated with goat anti-human aggrecan antibody (10 μg/ml, R&D Systems), goat anti-human Col2A1 antibody (4 μg/ml, Life Technologies, Paisley, UK) or normal goat IgG (control, 10 μg/ml) overnight at 2-8 °C. Following the overnight period, samples were washed with PBS (three times) and then incubated with a fluorescent secondary antibody – donkey anti-goat Alexa Fluor^®^ for one hour at room temperature (5 μg/ml, 594/488, Red/Green) (Life Technologies, Paisley UK). Control samples (MSCs grown on plastic under the same culture conditions) were also stained following the same protocol. Samples were imaged under a widefield microscope (Leica DMIRB inverted microscope, USA).

### Investigating Substrate Elasticity and Chondrogenic Signalling Pathways of MSC on Composite Substrates

To evaluate the cell morphological response to substrate elasticity, cells (on plastic and cultured on composite substrates) were fluorescently stained for paxillin (a focal adhesion protein) and their cytoskeleton (using a phallotoxin that binds to filamentous actin). Briefly, samples were washed with PBS (twice) then permeabilised at RT with 0.1 % Triton X-100 for ten minutes. Following a further wash step, the appropriate primary antibody was added; Anti-PXN rabbit antibody (5 μg/ml, Sigma Aldrich) or normal rabbit IgG control antibody (5 μg/ml, LifeTechnologies). Samples were then covered with foil and left overnight at 4 °C. Following the overnight period, samples were washed with PBS (twice). Secondary antibodies were then added to all samples; Alexa-Fluor 594 donkey anti-rabbit IgG (5 μg/ml, LifeTechnologies) and FITC-conjugated Phalloidin (2 μg/ml, Sigma Aldrich). Samples were covered with foil and left at room temperature for 1 hour. Samples were then washed with PBS (twice). A DAPI stain (DAPI solution, NucBlue^®^ Fixed Cell ReadyProbes™ Reagent, LifeTechnologies) was then added. Samples were covered with foil and left at room temperature for up to 15 minutes. The DAPI stain was then removed and samples washed with PBS (once). Samples were then left in PBS until imaging. Samples were imaged under a widefield microscope (Leica DMIRB inverted microscope, USA) at x10 magnification.

Blebbistatin (Sigma, UK), a nonmuscle myosin II (NMMII) inhibitor, was used to investigate the impact of substrate elasticity on stem cell differentiation potential. The toxicity and potency of the inhibitor when cultured with mesenchymal stem cells (MSCs) in culture was established in our lab and a concentration of 1 μM was used, in line with published values [22].

The toxicity and potency of the TGF-β type I receptor kinase inhibitor, SB505124 (Sigma,UK) when cultured with human MSCs was established [23]. A concentration of 1 μM was identified as least toxic with maximum inhibition (data not shown).

### Statistical Analysis

Bone marrow MSCs were used from four patients; *n=4* in all experiments. For dPCR, each cDNA sample was yielded from four biological repeats and were run in triplicate, with the average being used for gene expression calculations. Statistical significance between gene expression levels with and without the addition of inhibitors was assessed using a two-way ANOVA with Bonferroni post test. *p*≤0.05 was taken as significant. Data points on graphs show mean ± standard error of the mean (SE).

## Results & Discussion

### Human Stem Cell Chondrogenesis on Natural Polymer Composite Substrates

Composite substrates were prepared as discs of 8 mm diameter (Figure 1A) and characterised using scanning electron microscopy (SEM) and Fourier transform infrared spectroscopy (FTIR). SEM imaging was performed on film cross-sections showing no phase separation of polymer components in the substrates (Figure 1B). FTIR was carried out on all the raw processing components and confirmed the lack of modification to the characteristic amide groups of the silk component (1700-100 cm^-1^ amide I, 1600-1500 cm^-1^ amide II [24]) and the hydroxyl group of the cellulose component (around 1640 cm^-1^ adsorbed water [25]), thereby confirming the presence of both polymers in the cast composite substrate (Figure 1C).

**Figure 1.**
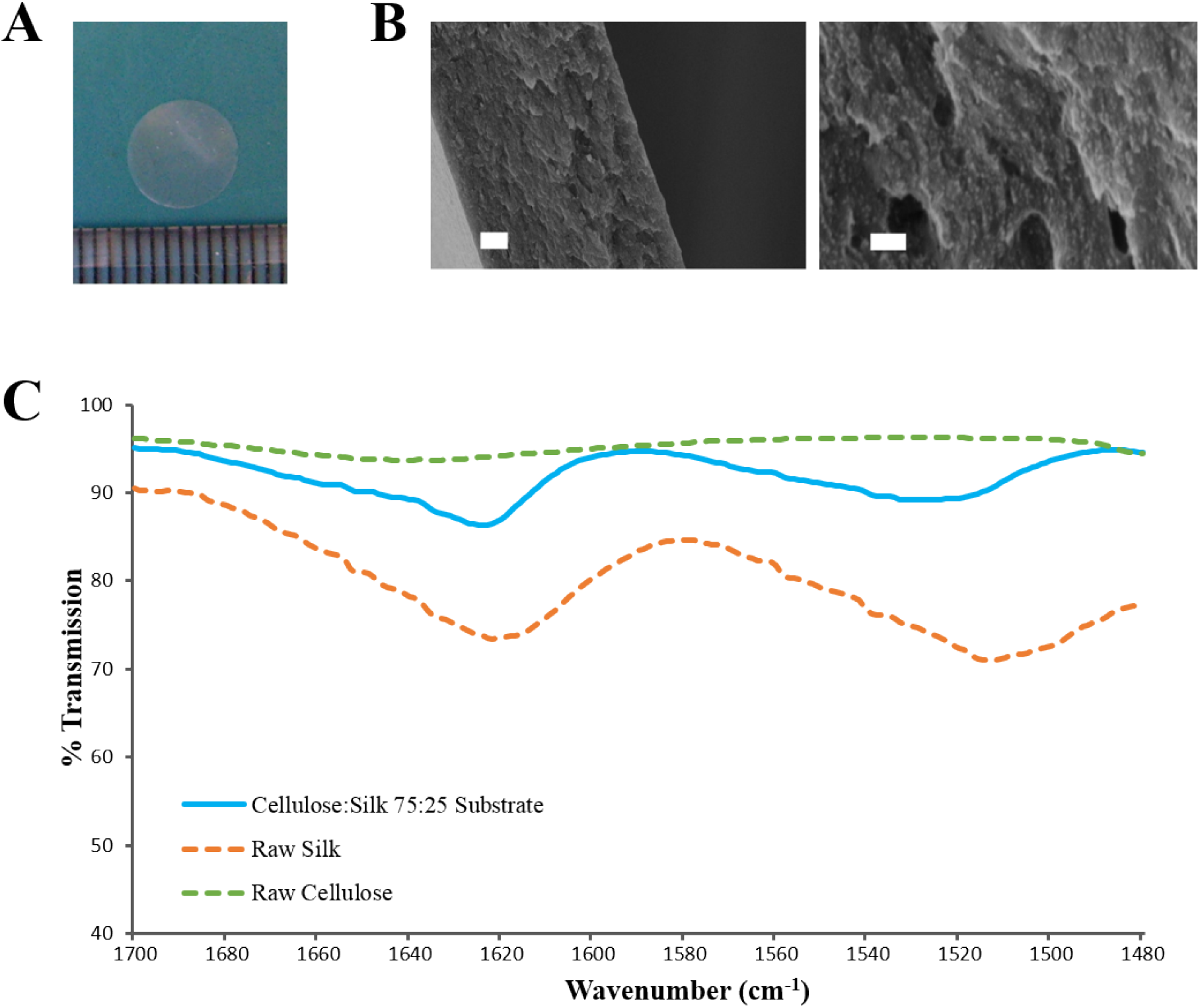
Composite Substrates of Cellulose and Silk. The natural polymers were blended in a 75:25 ratio and cast films produced following the protocol outlined by Singh *et al*. [17] A) Photo taken of 8mm disc of composite substrate, B) Scanning electron micrograph of film cross-section. Scale bars measure 2 μm (*left*) and 300 nm (*right*) respectively, C) FTIR was performed on the regenerated cellulose:silk composite substrates (solid line) and the raw cellulose and silk polymers (dashed lines).

The chondroinductive capacity of the composite substrate was tested using human bone marrow MSCs. Following culture, cells were analysed for chondrogenic gene expression and extracellular matrix protein deposition. MSCs were cultured on fibronectin coated composite substrates in stem cell expansion medium supplemented with 10 ng/ml FGF-2. This growth factor is known to enhance cellular proliferation and maintain stem cell differentiation potential [18] [19]. Cells were cultured for 14 days after which downstream gene analysis was performed. MSCs grown on these surfaces showed an upregulation in the chondrogenic genes type II collagen, aggrecan and Sox9, demonstrating the materials stimulatory capacity (Figure 2A). No upregulation in type I collagen (a marker of chondrocyte dedifferentiation) was seen. It should be noted that the levels of chondrogenic gene upregulation seen here were lower than those reported previously [17]. Some batch-to-batch variation when regenerating natural polymer composite substrates should be expected.

**Figure 2.**
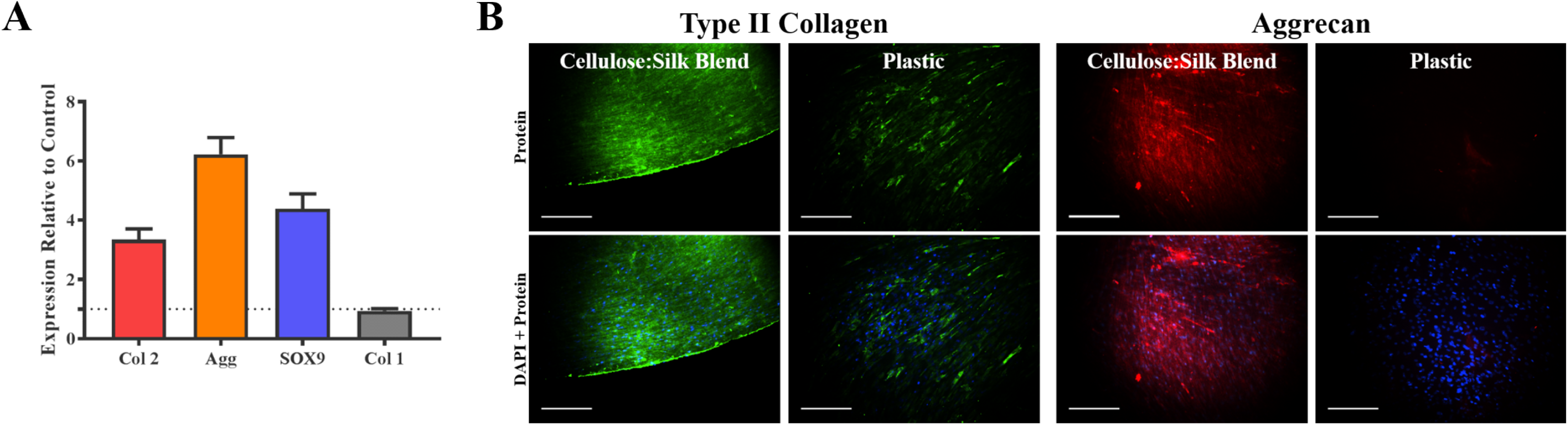
Mesenchymal Stem Cell Chondrogenesis on Composite Substrates. Human mesenchymal stem cells were cultured on fibronectin coated composite substrates in stem cell expansion medium supplemented with 10 ng/ml FGF-2. A) Cells were cultured for 14 days after which cells were screened for the expression of the chondrogenic genes - Type II collagen (Col2), Aggrecan (Agg), Sox9 and the marker for dedifferentiation Type I collagen (Col1). Gene expression levels were normalised to the expression of the *β*-actin housekeeping gene (shown in dotted line). Graph shows mean ± SE, n=4, B) Cells were also cultured for 21 days after which extracellular matrix (ECM) protein deposition was assessed for key cartilage proteins – Type II Collagen (green fluorescence) and Aggrecan (red fluorescence). Cell nuclei were stained using DAPI (blue). Representative images taken at x10 magnification, scale bar inset measures 300 μm.

The chondrogenic autoinductive potency of the composite substrate was demonstrated in the deposition of cartilage extracellular matrix (ECM) proteins (Figure 2B). Cells were cultured for 21 days after which ECM protein deposition was assessed using fluorescent antibody staining. Fluorescent regions arising from the deposition of the key earlier stage cartilage ECM proteins, type II collagen and aggrecan, were evident on the composite substrates, which were not present in control [26].

### The Chondrogenic Signalling Pathway and MSC Chondrogenesis on Composite Films

We hypothesised that the specific composition of cellulose and silk, in a 75:25 ratio, with its distinctive chemical functionalities (hydroxyl and amide respectively), is responsible for the stimulation of chondrogenesis in MSCs [17] [15]. To investigate this potential impact, we investigated the principal biochemical chondrogenic differentiation signalling pathway - TGFβ-driven Smad2/3 phosphorylation [8]. Phosphorylation of the Smad2/3 protein complex results in its translocation to the cell nucleus where it can regulate transcription factors and drive the upregulation of chondrogenic genes. Stimulation *via* this pathway leads to the upregulation of Sox9 which activates the expression of type II collagen genes [10]–[12]. This process occurs independently to substrate elasticity [27] and can be selectively inhibited using the small molecule inhibitor SB505124, a TGFβ type I receptor kinase inhibitor [28].

MSCs were seeded on the composite substrates in the presence of SB505124 and gene expression levels were assessed 14 days later (Figure 3). As a control, MSCs were seeded on plastic in the presence of TGFβ (Figure 3A). Addition of SB505124 to MSCs on plastic in the presence of TGFβ resulted in a significant down regulation in type II collagen and aggrecan genes with a reduction in Sox9 expression also noted; demonstrating the potency of the inhibitor (Figure 3A). When the inhibitor was added to composite substrate cultures, there was no statistical difference between treatment and control for the expression of the chondrogenic markers Sox9, aggrecan or type II collagen. The expression of type I collagen, a fibrochondrogenic marker, was also unchanged (Figure 3B). The lack of inhibition in MSC chondrogenesis on the natural polymer composites in the presence of SB505124 suggests the chondroinduction caused by the composite occurs independently to TGFβ Smad2/3 phosphorylation. Whilst this phosphorylation is regarded as the principal signalling mechanism for MSC chondrogenesis, it is important to consider other signalling pathways that may result in the same downstream effects on gene transcription. RhoA and Rho Kinase (ROCK) signalling can also stimulate the upregulation of Sox9 transcription resulting in the downstream expression of chondrogenic genes [12]. This pathway can be stimulated by TGFβ or mechanical stimulation [12] [29]. In the latter instance, conducive substrates have been shown to exert a specific pressure at which ROCK activity is optimal for cells to undergo chondrogenesis and results in autocrine TGFβ production [29]. This could suggest that the composite material is exerting the appropriate substrate stiffness to drive chondrogenic differentiation of the MSCs and would explain the lack of significant down regulation in chondrogenic gene expression (Figure 3B).

**Figure 3.**
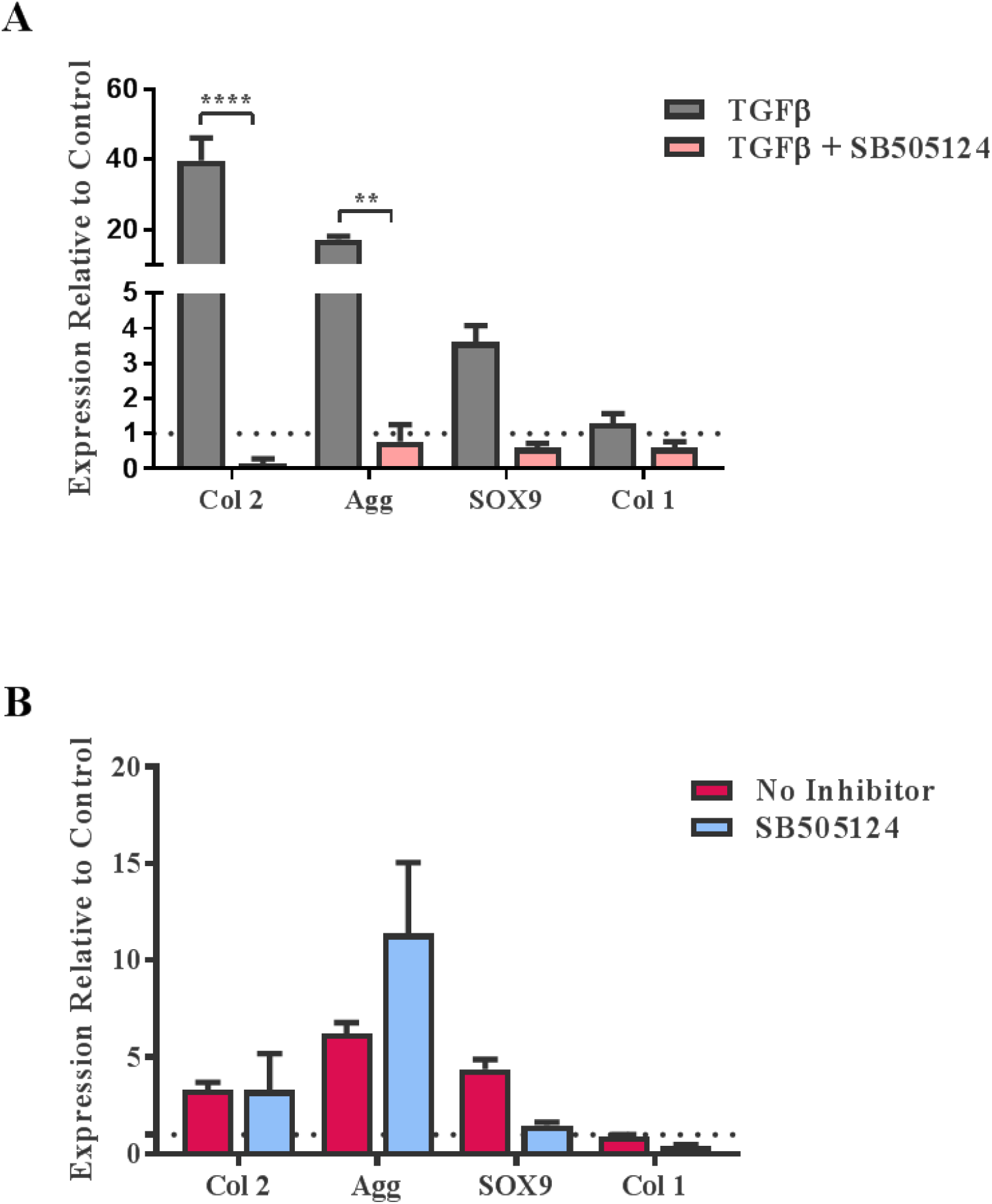
TGFβ – driven Smad2/3 Phosphorylation and Stem Cell Chondrogenesis. Human mesenchymal stem cells (MSCs) were cultured on A) Tissue culture plastic in chondrogenic media supplemented with transforming growth factor-β (TGF-β) and B) composite cellulose:silk 75:25 substrates in stem cell expansion media, with and without the SB505124 inhibitor. Cells were cultured for 14 days after which gene analysis was performed. Gene expression levels were normalised to expression levels of the housekeeping gene, β-actin (shown in dotted line). Statistical significance in B) was assessed using a Two-way ANOVA with Bonferroni post, *p*<0.05 taken as significant, data points show Mean ± SE, *n=4*.

### Substrate Elasticity and MSC Chondrogenesis on Composite Films

Singh *et al*. demonstrated that cellulose and silk blended in a 50:50 ratio and neat cellulose films, did not have the same chondroinductive effect on MSCs as cellulose-silk blended in a 75:25 ratio [17]. Further, this chondroinductive substrate exhibited stiffness and tensile strength intermediate to the former substrates [17]. Mammalian cells are known to respond to their substrate stiffness in both the number of adhesions formed to the surface and cell stretch [13] [30] and consequently in their differentiation [13]. Cells on stiff substrates form a greater number of adhesions with increased cytoskeletal elongation compared to cells on more elastic substrates.

To investigate whether the mechanical properties of the composite may influence MSC response to the substrate, the nature of MSC adhesion and morphology was investigated. MSCs seeded on plastic in the presence of biochemical chondroinductive medium (10 ng/ml TGFβ, Supplementary Info) or without TGFβ (Figure 4) were used as a reference. Cells were cultured under the appropriate media conditions with and without blebbistatin, a potent inhibitor of nonmuscle myosin II (NMM II) which is a component of the cell actin cytoskeleton important in force transmission between intra- and extra-cellular environments [14]. Following 3 days in culture, cells were stained for the actin cytoskeleton and the focal adhesion protein, paxillin.

**Figure 4.**
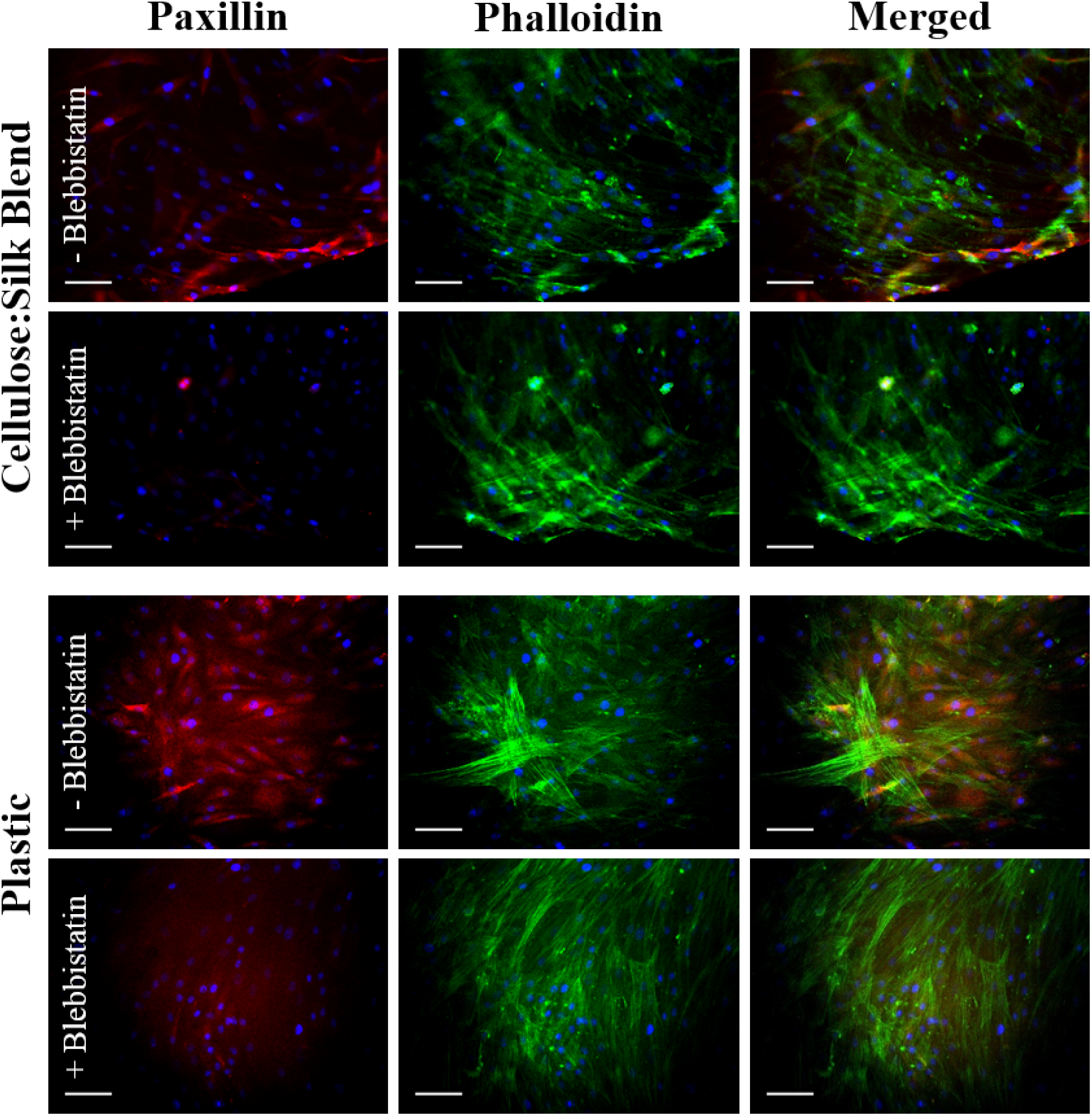
Stem Cell Adhesion and Morphology in Response to Substrate Elasticity. Human mesenchymal stem cells were cultured on cellulose:silk 75:25 composite substrates and tissue culture plastic in standard stem cell media. Following 3 days culture, cells were fluorescently stained for their focal adhesions (Paxillin, red) and cytoskeleton (Phalloidin, green) to assess the physical response of cells to their substrate with and without the use of Blebbistatin. Cell nuclei are stained using DAPI (blue). Images taken at x20 magnification, scale bar inset measures 100 μm. Representative images shown, *n*=4

Cells in chondroinductive medium grown on plastic had a polygonal, rounded shape, as expected of MSCs undergoing chondrogenic differentiation when stimulated with TGFβ (Supplementary info) [31]. The addition of blebbistatin reduced paxillin focal adhesion staining in all cases, suggesting a reduction in cell binding (Figures 4) [13] [30]. Cells grown on plastic in the absence of TGFβ showed elongated intense actin cytoskeletal structure and defined red fluorescent staining for the adhesion protein paxillin (Figure 4). Cells cultured on the composite substrate showed a lesser cell stretch with diffuse paxillin staining, demonstrating the MSC response to the elasticity of this surface (Figure 4). These cells however, did not demonstrate the distinct polygonal shape of MSCs undergoing chondrogenesis (Supplementary info), suggesting that the chondroinduction is partial and independent of the TGFβ signalling pathway. This finding further supports the results seen in the presence of SB505124 (Figure 3) and suggests that the potential mechanism for the chondroinduction of MSCs grown on the cellulose-silk composites is dependent on surface elasticity.

To evaluate the level of elasticity-driven chondroinduction on these natural polymer composites, chondrogenic gene expression levels were investigated for MSCs grown on the substrates in the presence and absence of blebbistatin (Figure 5). Inhibition of the NMM II cytoskeletal protein by blebbistatin resulted in a significant downregulation in the expression of chondrogenic markers (Figure 5). The expression of type I collagen, a marker of undifferentiated MSCs, was not affected. Inhibition of the NMM II cytoskeletal protein complex has been specifically implicated in preventing substrate elasticity driven differentiation in MSCs [13] and our results support this finding. It is worth noting that blebbistatin did not fully downregulate the chondrogenic markers to basal levels of the untreated control. This suggests that other signalling pathways that mediate cellular mechanotransduction may be implicated. The mechanical properties of this composite may support the induction of autocrine TGFβ production via ROCK signalling in conjunction with the NMM II mediated differentiation [29].

**Figure 5.**
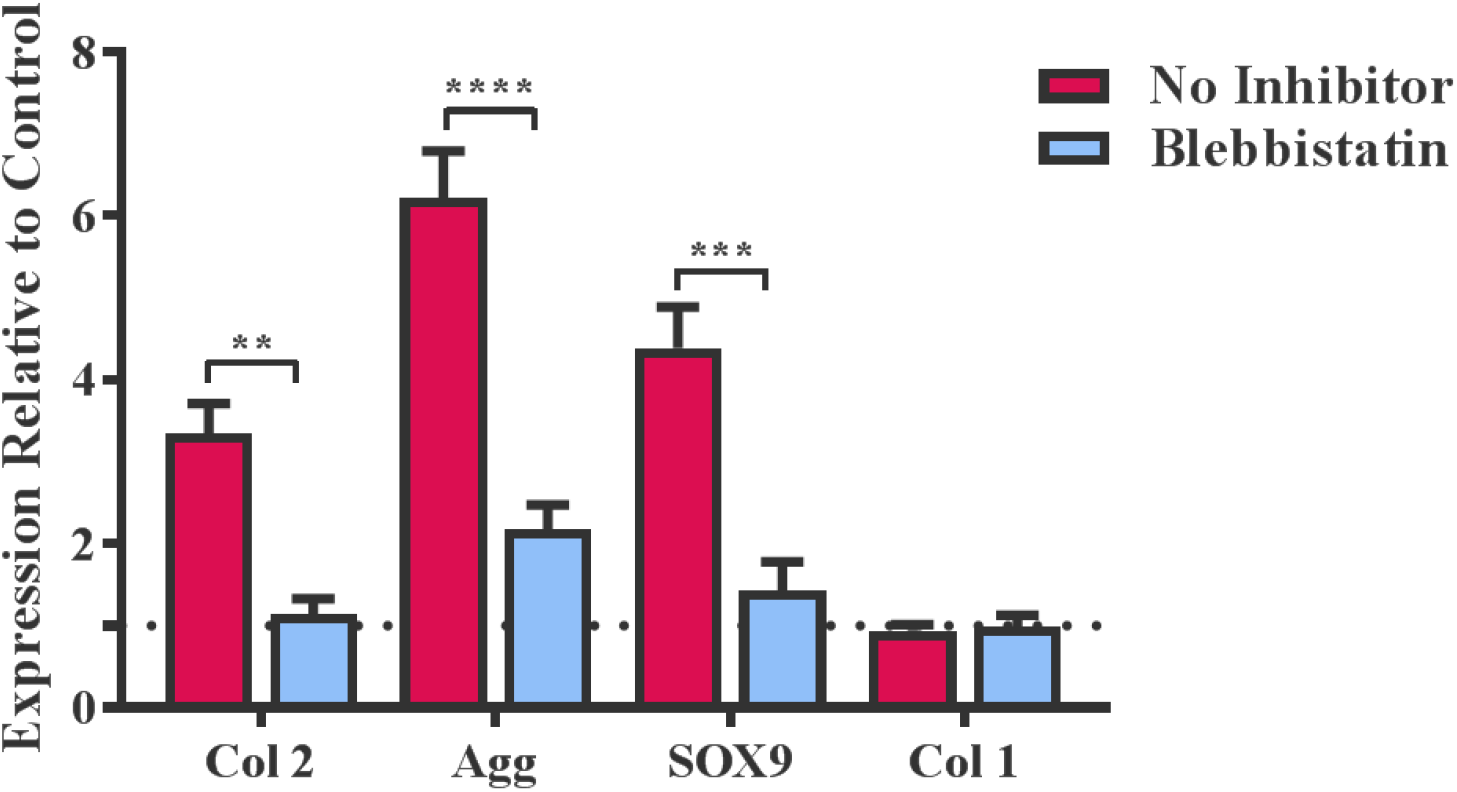
Substrate Elasticity and Stem Cell Chondrogenesis. Human mesenchymal stem cells (MSCs) were cultured on composite cellulose:silk 75:25 substrates in stem cell expansion media, with and without the Blebbistatin inhibitor. Cells were cultured for 14 days after which gene analysis was performed. Gene expression levels were normalised to expression levels of the housekeeping gene, *β*-actin (shown in dotted line). Significant differences in gene expression was assessed using a Two-way ANOVA with Bonferroni post-test, p<0.05 taken as significant. Data points show Mean ± SE, *n=4*.

## Conclusion

Potential mechanisms for the chondroinduction of MSCs cultured on cellulose:silk 75:25 composite substrates were investigated. The chondroinduction did not involve the principal chondrogenic differentiation signalling pathway - Smad2/3 regulated phosphorylation *via* TGF-β signalling. However, inhibition of force transmission between intracellular and extracellular environments *via* NMM II significantly reduced chondrogenic gene expression of MSCs on these substrates. Further investigations are needed to reveal the interoperation of signalling pathways that orchestrate the transduction of physical cues into biochemical cues to mediate this differentiation pathway.

Harnessing the mechanical properties of this natural polymer in a 3D construct with the appropriate porosity and structural integrity will enable cartilage tissue engineering, without the need for stimulating growth factors, thereby supporting the development of a smart biomaterial. The inherent chondroinductive capacity of these natural polymer composites can facilitate the recruitment of MSCs at the implantation site [32], proposing a nascent cell population for enhancing the materials regenerative impact.

## Acknowledgements

The authors would like to thank the EPSRC (Doctoral Training Partnership, EP/K502996/1) for funding this work.

## Supplementary Info

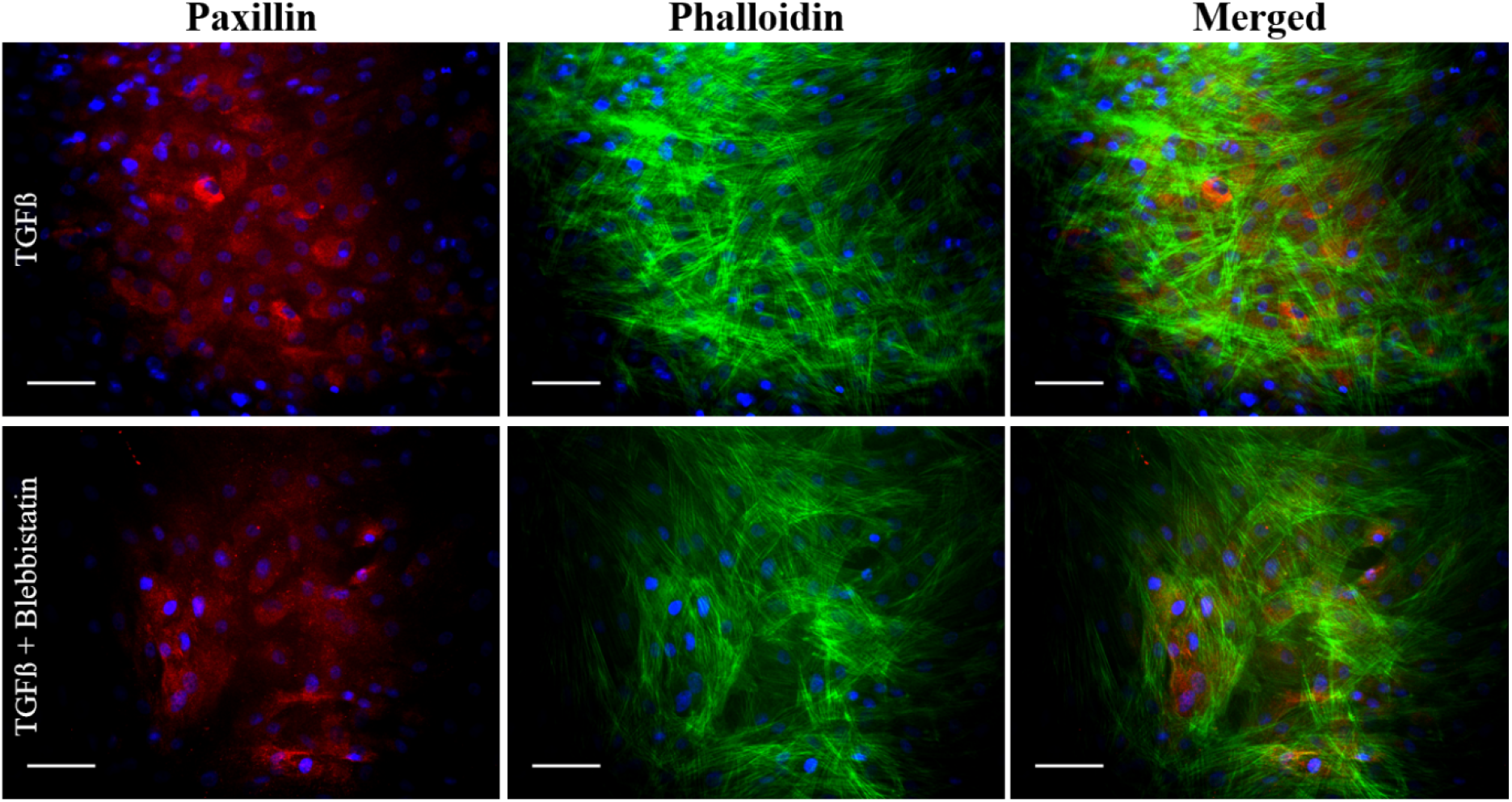
Stem Cell Adhesion and Morphology in Response to Stimulating Factors. Human mesenchymal stem cells were cultured on plastic in the presence of the chondrogenic stimulator transforming growth factor-β (TGF-β). Following 3 days culture, cells were fluorescently stained for their focal adhesions (Paxillin, red) and cytoskeleton (Phalloidin, green) to assess the physical response of cells undergoing chondrogenesis under TGF-β stimulation with and without the use of Blebbistatin. Cell nuclei are stained using DAPI (blue). Images taken at x20 magnification, scale bar inset measures 100um. Representative images shown, *n*=4

## References

[1] W. Kafienah, S. Mistry, S. C. Dickinson, T. J. Sims, I. Learmonth, and A. P. Hollander, “Three-dimensional cartilage tissue engineering using adult stem cells from osteoarthritis patients,” Arthritis Rheum., vol. 56, no. 1, pp. 177–187, 2007.

[2] H. J. Lee, J. S. Lee, T. Chansakul, C. Yu, J. H. Elisseeff, and S. M. Yu, “Collagen mimetic peptide-conjugated photopolymerizable PEG hydrogel,” Biomaterials, vol. 27, no. 30, pp. 5268–5276, 2006.

[3] L. Meinel, S. Hofmann, V. Karageorgiou, L. Zichner, R. Langer, D. Kaplan, and G. Vunjak-Novakovic, “Engineering cartilage-like tissue using human mesenchymal stem cells and silk protein scaffolds,” Biotechnol. Bioeng., vol. 88, no. 3, pp. 379–391, 2004.

[4] R. M. Coleman, N. D. Case, and R. E. Guldberg, “Hydrogel effects on bone marrow stromal cell response to chondrogenic growth factors,” Biomaterials, vol. 28, no. 12, pp. 2077–2086, 2007.

[5] H. M. Van Beuningen, H. L. Glansbeek, P. M. Van Der Kraan, and W. B. Van Den Berg, “Osteoarthritis-like changes in the murine knee joint resulting from intra-articular transforming growth factor-b injections,” Osteoarthr. Cartil., vol. 8, no. 1, pp. 25–33, 2000.

[6] D. T. Scadden, “The stem-cell niche as an entity of action.,” Nature, vol. 441, no. 7097, pp. 1075–1079, 2006.

[7] S. Martino, F. D’ Angelo, I. Armentano, J. M. Kenny, and A. Orlacchio, “Stem cell-biomaterial interactions for regenerative medicine,” Biotechnol. Adv., vol. 30, no. 1, pp. 338–351, 2012.

[8] B. Johnstone, T. M. Hering, a I. Caplan, V. M. Goldberg, and J. U. Yoo, “In vitro chondrogenesis of bone marrow-derived mesenchymal progenitor cells.,” Exp. Cell Res., vol. 238, no. 1, pp. 265–272, 1998.

[9] J. M. Yingling, K. L. Blanchard, and J. S. Sawyer, “Development of TGF-beta signalling inhibitors for cancer therapy.,” Nat. Rev. Drug Discov., vol. 3, no. 12, pp. 1011–1022, 2004.

[10] T. Furumatsu, M. Tsuda, N. Taniguchi, Y. Tajima, and H. Asahara, “Smad3 induces chondrogenesis through the activation of SOX9 via CREB-binding protein/p300 recruitment,” J. Biol. Chem., vol. 280, no. 9, pp. 8343–8350, 2005.

[11] Y. Kamachi, M. Uchikawa, and H. Kondoh, “Pairing SOX off: With partners in the regulation of embryonic development,” Trends Genet., vol. 16, no. 4, pp. 182–187, 2000.

[12] D. R. Haudenschild, J. Chen, N. Pang, M. K. Lotz, and D. D. D’Lima, “Rho kinase-dependent activation of SOX9 in chondrocytes,” Arthritis Rheum., vol. 62, no. 1, pp. 191–200, 2010.

[13] A. J. Engler, S. Sen, H. L. Sweeney, and D. E. Discher, “Matrix Elasticity Directs Stem Cell Lineage Specification,” Cell, vol. 126, no. 4, pp. 677–689, 2006.

[14] A. F. Straight, A. Cheung, J. Limouze, I. Chen, N. J. Westwood, J. R. Sellers, and T. J. Mitchison, “Dissecting temporal and spatial control of cytokinesis with a myosin II Inhibitor.,” Science, vol. 299, no. 5613, pp. 1743–1747, 2003.

[15] J. M. Curran, R. Chen, and J. a. Hunt, “Controlling the phenotype and function of mesenchymal stem cells in vitro by adhesion to silane-modified clean glass surfaces,” Biomaterials, vol. 26, no. 34, pp. 7057–7067, 2005.

[16] M. J. Dalby, N. Gadegaard, R. Tare, A. Andar, M. O. Riehle, P. Herzyk, C. D. W. Wilkinson, and R. O. C. Oreffo, “The control of human mesenchymal cell differentiation using nanoscale symmetry and disorder.,” Nat. Mater., vol. 6, no. 12, pp. 997–1003, 2007.

[17] N. Singh, S. S. Rahatekar, K. K. K. Koziol, T. S. Ng, A. J. Patil, S. Mann, A. P. Hollander, and W. Kafienah, “Directing chondrogenesis of stem cells with specific blends of cellulose and silk,” Biomacromolecules, vol. 14, no. 5, pp. 1287–1298, 2013.

[18] I. Martin, A. Muraglia, G. Campanile, R. Cancedda, and R. Quarto, “Fibroblast Growth Factor-2 Supports ex Vivo Expansion and Maintenance of Osteogenic Precursors from Human Bone Marrow 1,” Endocrinology, vol. 138, no. 10, pp. 4456–4462, 1997.

[19] L. a. Solchaga, K. Penick, J. D. Porter, V. M. Goldberg, A. I. Caplan, and J. F. Welter, “FGF-2 enhances the mitotic and chondrogenic potentials of human adult bone marrow-derived mesenchymal stem cells,” J. Cell. Physiol., vol. 203, no. 2, pp. 398–409, 2005.

[20] E. Ruoslahti, “Fibronectin in cell adhesion and invasion,” Cancer Metastasis Rev., vol. 3, no. 1, pp. 43–51, 1984.

[21] H. M. Kowalczyn, M. Nowak-wyrzykowska, J. Dobkowski, and R. Kołos, “Adsorption characteristics of human plasma fibronectin in relationship to cell adhesion,” 2001.

[22] R. K. Das, O. F. Zouani, C. Labrugère, R. Oda, and M. C. Durrieu, “Influence of nanohelical shape and periodicity on stem cell fate,” ACS Nano, vol. 7, no. 4, pp. 3351–3361, 2013.

[23] Y. Tang, X. Wu, W. Lei, L. Pang, C. Wan, Z. Shi, L. Zhao, T. R. Nagy, X. Peng, J. Hu, X. Feng, W. Van Hul, M. Wan, and X. Cao, “TGF-beta1-induced migration of bone mesenchymal stem cells couples bone resorption with formation.,” Nat. Med., vol. 15, no. 7, pp. 757–765, 2009.

[24] A. Barth and C. Zscherp, “What vibrations tell about proteins,” Q. Rev. Biophys., vol. 35, no. 4, p. S0033583502003815, 2002.

[25] P. Garside and P. Wyeth, “Identification of Cellulosic Fibres by FTIR Spectroscopy,” Stud. Conserv., vol. 48, no. 4, pp. 269–275, 2003.

[26] J. Xu, W. Wang, M. Ludeman, K. Cheng, T. Hayami, J. C. Lotz, and S. Kapila, “Chondrogenic Differentiation of Human Mesenchymal Stem Cells in Three-Dimensional Alginate Gels,” Tissue Eng. Part A, vol. 14, no. 5, pp. 667–680, 2008.

[27] J. S. Park, J. S. Chu, A. D. Tsou, R. Diop, Z. Tang, A. Wang, and S. Li, “The effect of matrix stiffness on the differentiation of mesenchymal stem cells in response to TGF-β,” Biomaterials, vol. 32, no. 16, pp. 3921–3930, 2011.

[28] S. DaCosta Byfield, C. Major, N. J. Laping, and A. B. Roberts, “SB-505124 is a selective inhibitor of transforming growth factor-beta type I receptors ALK4, ALK5, and ALK7.,” Mol. Pharmacol., vol. 65, no. 3, pp. 744–752, 2004.

[29] J. L. Allen, M. E. Cooke, and T. Alliston, “ECM stiffness primes the TGF pathway to promote chondrocyte differentiation,” Mol. Biol. Cell, vol. 23, no. 18, pp. 3731–3742, 2012.

[30] Y.-L. Pelham, Robert J Wang, “Cell locomotion and focal adhesions are regulated by substrate flexibility,” Proc. Am. Thorac. Soc., vol. 94, pp. 13661–13665, 1997.

[31] L. Gao, R. McBeath, and C. S. Chen, “Stem cell shape regulates a chondrogenic versus myogenic fate through rac1 and N-cadherin,” Stem Cells, vol. 28, no. 3, pp. 564–572, 2010.

[32] a. Nair, J. Shen, P. Lotfi, C. Y. Ko, C. C. Zhang, and L. Tang, “Biomaterial implants mediate autologous stem cell recruitment in mice,” Acta Biomater., vol. 7, no. 11, pp. 3887–3895, 2011.

